# Novel ionic liquids-based extraction method that preserves molecular structure from cutin

**DOI:** 10.1101/2020.06.01.127837

**Authors:** Carlos J.S. Moreira, Artur Bento, Joana Pais, Johann Petit, Rita Escórcio, Vanessa G. Correia, Ângela Pinheiro, Łukasz P. Haliński, Oleksandr O. Mykhaylyk, Christophe Rothan, Cristina Silva Pereira

## Abstract

The biopolyester cutin is ubiquitous in land plants, building the polymeric matrix of the plant’s outermost defensive barrier - the cuticle. Cutin influences many biological processes *in planta* however due to its complexity and highly branched nature, the native structure remains partially unresolved. Our aim was to define an original workflow for the purification and systematic characterisation of the molecular structure of cutin. To purify cutin we tested the ionic liquids cholinium hexanoate and 1-butyl-3-methyl-imidazolium acetate. The ensuing polymers are highly esterified, amorphous and have the typical monomeric composition as demonstrated by solid state NMR, complemented by spectroscopic (GC-MS), thermal (DSC) and x-ray scattering (WAXS) analyses. A systematic study by solution-state NMR of cryogenically milled cutins extracted from Micro-Tom tomatoes (the wild type and the *gpat6* and *cus1* mutants) was undertaken. Their molecular structures, relative distribution of ester aliphatics, free acid end-groups and free hydroxyl groups, differentiating between those derived from primary and secondary esters, were solved. The acquired data demonstrate the existence of free hydroxyl groups in cutin and reveal novel insights on how the mutations impact the esterification arrangement of cutin. Compared to conventional approaches, the usage of ionic liquids for the study of plant polyesters opens new avenues since simple modifications can be applied to recover a biopolymer carrying distinct types/degrees of modifications (*e.g*. preservation of esters or cuticular polysaccharides), which in combination with the solution NMR methodologies developed here, constitutes now essential tools to fingerprint the multi-functionality and the structure of cutin *in planta*.

## Introduction

Plant polyesters, namely cutin and suberin, are the third most abundant plant polymers right after cellulose/hemicellulose and lignin. Naturally, due to their high abundance in nature, plant polyesters are considered as promising substitutes to petroleum-based plastics (Heredia-Guerrero et al., 2017). In particular, cutin makes up the polymeric matrix of the cuticle that builds the protective layer of the aerial parts of land plants; an evolutionary feature acquired during the colonization of terrestrial environments (Fich et al., 2016). The cuticle constituents (cutin and waxes) are deposited onto the polysaccharide layer of the walls of the epidermal cells (Segado et al., 2016). Cutin is, in general, a highly branched polymer, mainly composed of C16 and C18 fatty acids, containing mostly terminal (ω-hydroxyl) and *mid*-chain hydroxyl group, linked through ester bonds. Other functional groups such as aromatics, dicarboxylic acids and glycerol can also be found in cutin at low amounts (Mazurek et al., 2017).

Over the years, many authors have contributed to elucidate the roles played by cutin in diverse biological processes along plant development, growth and response to biotic and/or abiotic stresses (Fich et al., 2016). However, current methods for the extraction and analysis of cutin polyesters have inherent limitations. The extraction of cutin from a plant source usually relies on time-consuming processes that include enzymatic digestion of polysaccharides followed by thorough organic solvent extraction of the soluble waxes present in the cuticle (Chatterjee et al., 2012). In addition, the most frequent chemical analysis of cutin are based on total/partial hydrolyses of the polyesters and therefore disclose solely the monomeric constituents attained through (Graça and Lamosa, 2010; Fernández et al., 2016), regardless that sometimes solid state spectroscopic based analyses of the polymer are also used (Deshmukh et al., 2003; Fernández et al., 2016). The monomeric constituents can disclose a partial view of the basic composition of the biopolymer (*i.e.* of the hydrolysable constituents), while providing insights on its biosynthesis (Bakan and Marion, 2017) but not of their supra-molecular organisation, which remains largely unknown (Fich et al., 2016; Bakan and Marion, 2017). To advance our understanding of important cutin-related questions such as cutin/cell wall polymers interactions or the role of cutin in defence to pathogens (Chatterjee et al., 2016), a better insight into the structure of cutin in its native state is highly required.

Ionic liquids – usually defined as salts on a liquid state below 100 °C – may facilitate the processing of plant polymers due to their capacity to induce swelling/solubilisation and/or to catalyse the cleavage of specific inter-molecular bonds (Rogers and Seddon, 2003). In particular, some imidazolium-based ionic liquids can efficiently disrupt the intermolecular hydrogen bonding between hydroxyl groups in cellulose (Li et al., 2018) whereas some cholinium alkanoates can catalyse selectively the hydrolysis of inter-molecular acylglycerol esters (Garcia et al., 2010; Ferreira et al., 2012; Ferreira et al., 2014). The latter ionic liquid was used by us to extract suberin from cork - a plant polyester sharing chemical similarities with cutin – by catalysing a selective and mild hydrolysis of acyl glycerol esters yet preserving most extant linear aliphatic esters (Ferreira et al., 2014; Correia et al., 2020).

Our aim was to establish a novel cutin extraction method that allows the study of native cutin architecture and properties and is applicable to a wide range of plant species and tissues. To meet these criteria, the newly-developed method should be easy to process and rapid and should preserve the chemical structure of the cutin. To this end, we first established an ionic liquid approach for the extraction of cutin, with the solubilisation of cutin from tomato peel as a proof-of-concept. We demonstrated that the cholinium hexanoate process renders a near native cutin as shown by Scanning Electronic Microscopy (SEM), ^13^C Magic Angle Spinning Nuclear Magnetic Resonance (^13^C MAS NMR), and Differential Scanning Calorimetry (DSC) analyses of the cutin structure. These analyses were complemented by Gas Chromatography – Mass Spectrometry (GC-MS) analyses of the hydrolysable constituents. In addition, we established for the first time the molecular structure in solution of near native cutins (solubilised with the aid of cryogenic milling) through high-resolution one- and two-dimensional solution state NMR analyses. Extension of our approach from a processing tomato cultivar to the miniature Micro-Tom cultivar, including two cutin biosynthesis and polymerisation mutants, highlighted the consistency of our findings with published results but also revealed new features of near native cutin. We therefore believe that our methodological approach will support discovery in the field of cutin biogenesis and biosynthesis.

## Results

### A highly esterified cutin was purified using ionic liquids that mediate mostly the dissolution of sub-cuticular polysaccharides

Seeking to establish a novel methodology to extract cutin from tomato peels, we resorted to cholinium hexanoate and 1-butyl-3-methyl-imidazolium acetate (hereafter defined as BMIM acetate). Cholinium hexanoate was chosen because of its ability to mediate the extraction of suberin from cork through mild and selective hydrolysis of acylglycerol ester bonds (Ferreira et al., 2014), and BMIM acetate due to its proven ability to mediate the dissolution of cellulose (Li et al., 2018). First, we tested if either ionic liquid (100 °C without stirring) hydrolyses glyceryl trioctanoate and octyl octanoate that contain an acylglycerol ester bond and a linear aliphatic ester bond, respectively. Glyceryl trioctanoate was hydrolysed in the presence of both ionic liquids, yet the efficiency of the reaction was higher when cholinium hexanoate was used (Fig. 1). Cholinium hexanoate did not catalyse the cleavage of octyl octanoate (Fig. 1A), contrary to the BMIM acetate that catalysed this reaction though inefficiently (Fig. 1B). As previously reported, cholinium hexanoate catalyses specifically the hydrolysis of acylglycerol esters (Ferreira et al., 2014), regardless that in the present study the absence of agitation and the higher water content of the ionic liquid reduced the reaction efficiency.

**Fig. 1.**
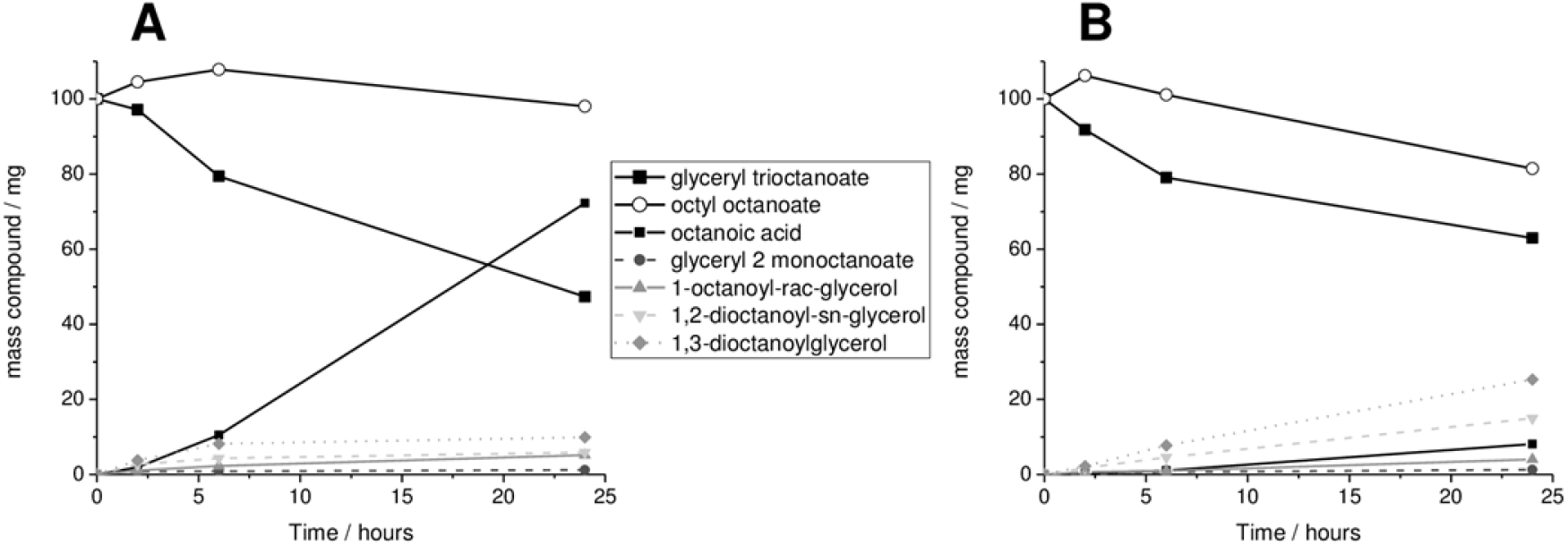
Compounds detected after the reaction of glyceryl trioctanoate and octyl octanoate with either cholinium hexanoate (A) or 1-butyl-3-methylimidazolium acetate (B) for 2, 6 and 24 hours (the observed average standard errors were negligible, < 4%). All compounds were identified and quantified by GC-MS. At time zero, glyceryl trioctanoate and octyl octanoate were assumed to represent the only compounds present in mixture.

We then tested the potential of these ionic liquids for the isolation of cutin from tomato peels after 2, 15 and 170 hours compared to a conventional method (*i.e.* enzymatic removal of polysaccharides followed by organic solvent mediated dewaxing). In the process of suberin extraction from cork using cholinium hexanoate, suberin in the filtrate is recovered by precipitation in an excess of water (Ferreira et al., 2012). We preliminarily tested a 2 hours reaction of cutin peels in cholinium hexanoate, and verified using ATR-FTIR that the archetypal bands assigned to cutin, *i.e.* long chain aliphatics (CH_2_ and C=O), were detected in the insoluble fraction (not in the filtrate as observed for cork suberin) whereas the filtrate shows enrichment in bands usually assigned to polysaccharides (C-O-C) (Supplementary Fig. S1). Accordingly, the produced insoluble fractions were characterised using SEM (Fig. 2) and ^13^C MAS NMR (Fig. 3A). SEM imaging of the cutins extracted with either ionic liquid are virtually identical: a clean thick cutin-continuum showing the epidermal cells grooves (Fig. 2A-F). In the reference cutin, *i.e.* obtained through the conventional enzymatic-based process, the cutin-continuum apparently overlaps with other cellular components, and many intracellular spaces are not hollow (Fig. 2G).

**Fig. 2.**
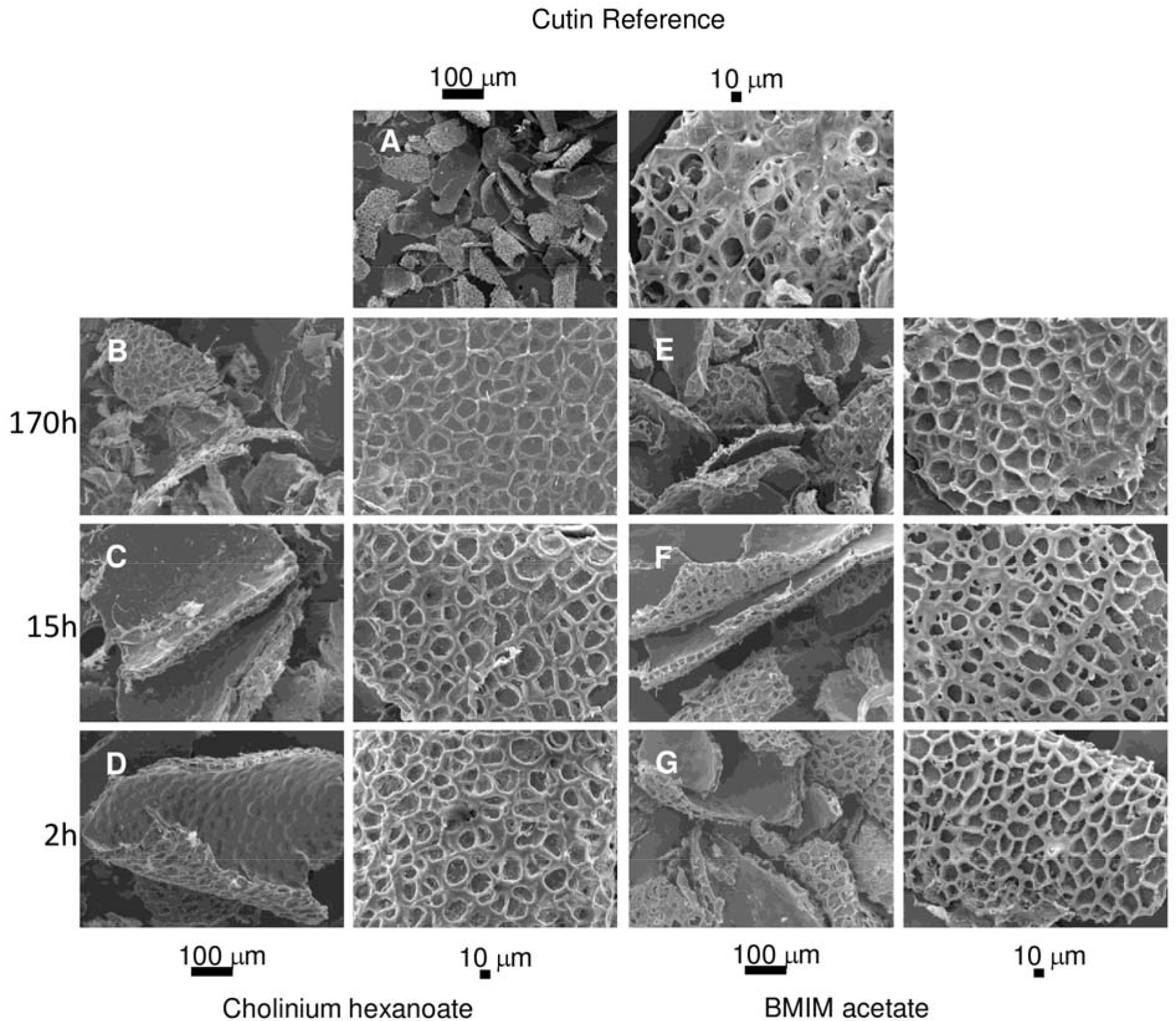
SEM imaging of cutin purified after treatment with cholinium hexanoate (B-D) or 1-butyl-3-methylimidazolium acetate (E-G) after 2, 15 and 170 hours. All samples show a clean thick cutin-continuum comprising the epidermal cells grooves. A representative cutin reference sample (*i.e.* obtained through the conventional enzymatic-based process) is also shown denoting many intracellular spaces that are not hollow (A).

**Fig. 3.**
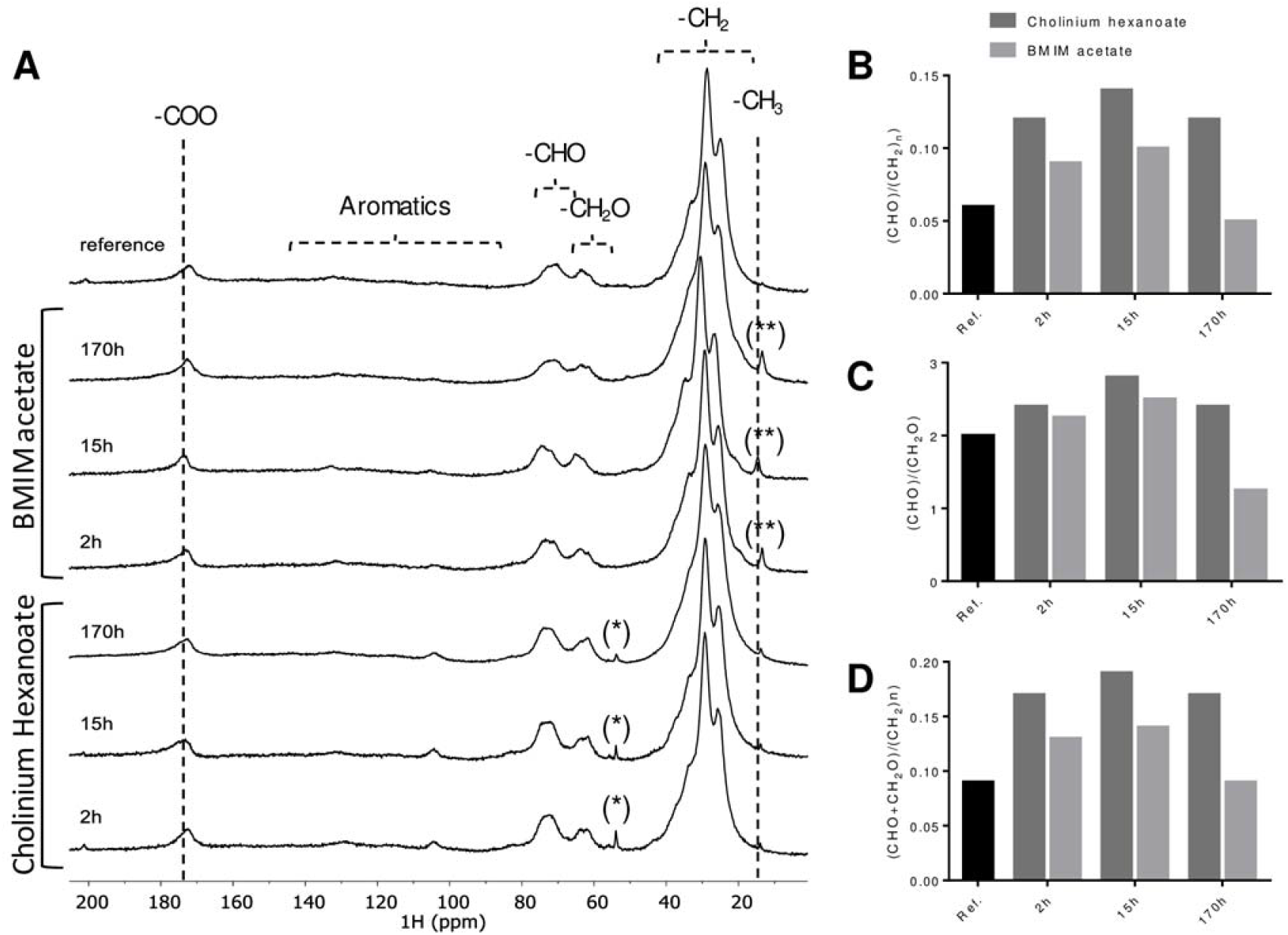
^13^C MAS NMR spectra obtained for the cutin reference and the cutin samples derived from reactions with cholinium hexanoate or 1-butyl-3-methylimidazolium acetate after 2, 15 and 170 hours (A) and the corresponding calculated reticulation (B-C) and esterification (D) ratios. The regions assigned to the long methylene chains, the oxygenated aliphatics, aromatics and the carboxyl groups are marked. The imidazolium-based cation contributes to the signal assigned to the CH_3_ groups (15 ppm **), whereas the cholinium cation is seen in the signal at 54 ppm *; both contaminants can be washed out.

In the ^13^C MAS NMR spectrum of the reference cutin, the major structural classes assigned to cutin include the long methylene chains - (CH_2_)_n_ with major peaks at 26, 29 & 34 ppm, the oxygenated aliphatics - CH_2_O (63 ppm) and CHO (73 ppm), and the carboxyl groups at 172 ppm, comprising the contribution of both esters and acids (Chatterjee et al., 2016) (Fig. 3A). Only minor signals can be assigned to the aromatic region (105 & 130 ppm). The spectral signatures of the remaining cutins are very similar regardless of used ionic liquid and extraction time and also similar to the reference cutin spectrum (Fig. 3 and Table 1). The relative contributions of the signals assigned to aromatics for the cutins purified with either ionic liquid increased along the reaction time, possibly an artefact derived from phase corrections. The relative contributions of the oxygenated aliphatics region (57-92 ppm) are higher in the ionic liquid extracted cutins compared to the reference cutin (Fig. 3A). This region might also comprise resonances derived from polysaccharides (Chatterjee et al., 2016). Most subcuticular polysaccharides can be removed from cutin using an acidic hydrolysis mediated by TFA (Arrieta-Baez and Stark, 2006), regardless that cellulose might not be totally removed (Hernández Velasco et al., 2017). In the present study, the NMR spectra of cutins obtained using the cholinium hexanoate (2 h reaction) before and after the acidic hydrolysis treatment are virtually identical (Table 1, Supplementary Fig. S2). The few observed alterations can be explained by the hydrolysis of esters during the acidic treatment (Arrieta-Baez and Stark, 2006). Based on these results, the oxygenated aliphatics region can be mostly assigned to cutin. Consequently, the biopolymer reticulation level can be reasonably estimated through the ratio of signal’s integral in the CHO region of the oxygenated aliphatics (67-92 ppm) with that of the entire aliphatic region (8-50 ppm) or that of the CH_2_O region (57-67 ppm) (Matas et al., 2011; Chatterjee et al., 2016). Based on the calculated reticulation ratios, cholinium hexanoate usage apparently rendered a biopolymer displaying higher reticulation compared to either that attained with the BMIM acetate or the conventional approach (Fig. 3B-C). At this stage, one cannot exclude that the presence of cellulose embedded in the biopolymer might increase the estimated reticulation levels. A similar trend was observed when estimating their esterification levels (Fig. 3D), which can be inferred through the ratio between the integral of the total oxygenated aliphatic region (CHO & CH_2_O) with that of the entire aliphatic region (8-50 ppm) (Matas et al., 2011). The esterification of the cholinium cation with cutin’s free acids was reported before as mechanistically very unlikely (Ferreira et al., 2014).

**Table 1.** Relative abundance of the contributions of the region of the aliphatics (10-50 ppm), oxygenated aliphatics (57-92 ppm), aromatics (92-165 ppm) and carboxyl groups (165-185 ppm) for each cutin ^13^C MAS NMR spectrum.

| Method | Cutin major structural classes | Relative abundance of the $^{13}\text{C}$ MAS NMR assigned regions (%) | | | | |
| --- | --- | --- | --- | --- | --- | --- |
| | | $\text{C}=\text{O}$ | $\text{C}=\text{C}$ | $\text{CHO}$ | $\text{CH}_2\text{O}$ | $(\text{CH}_2)_n$ |
|  |  | carboxyl | aromatics | oxygenated | aliphatics | aliphatics |
| Reference | - | 5.1 | 1.7 | 5.1 | 2.6 | 85.5 |
| Cholinium hexanoate | 2 h | 4.8 | 1.6 | 9.6 | 3.7 | 80.0 |
|  | 2 h + TFA | 3.9 | 3.1 | 10.2 | 4.7 | 78.1 |
|  | 15 h | 4.7 | 3.1 | 10.9 | 3.9 | 77.5 |
|  | 170 h | 4.7 | 3.1 | 9.1 | 3.9 | 65.0 |
| Imidazolium acetate | 2 h | 5.6 | 4.0 | 7.2 | 3.2 | 80.0 |
|  | 15 h | 5.5 | 5.5 | 7.8 | 3.1 | 73.5 |
|  | 170 h | 5.6 | 6.5 | 4.0 | 3.2 | 80.6 |

In order to further elucidate if the ionic liquid-based extractions can indeed render a near native-cutin we resorted to DSC (Fig. 4A) and WAXS (Fig. 4B) measurements of the cutins extracted with the ionic liquids (2 and 170 hours) together with the reference cutin. The DSC thermograms are depicted in Fig. 4A. The cutins extracted using either ionic liquid for 2 hours show higher enthalpy energies for melting the biopolymer and lower melting temperatures (ΔH=129.6 J·g^−1^ & T_m_ = 100.5 °C and ΔH=97.5 J·g^−1^ & T_m_ = 100.6°C, for cholinium hexanoate and BMIM acetate, respectively) compared to the reference cutin (ΔH=70.5 J·g^−1^ & T_m_ = 114.5 °C). All thermograms show a relatively broad melting curve (Fig. 4A), which is typical for heterogeneous and amorphous materials (Benítez et al., 2018) and similar glass transition temperatures (ca. −20 °C). The peaks for the cutins which originated from the 2 hours ionic liquid reactions are less broad compared to the reference cutin, suggestive of increased homogeneity. This feature was lost when extensive reaction times were used, consistent with the estimated reduction in the biopolymer reticulation and esterification (Fig. 3B-D). The WAXS patterns of all cutin samples (Fig. 4B) are mainly represented by a broad diffuse peak with the maximum intensity at *q* ~ 1.41 Å^−1^ which most likely corresponds to an amorphous structure commonly formed by organic polymeric materials with an inter-chain distance of around 4.5 Å. Considering cutin’s composition, this amorphous structure should be related to randomly-packed acyl chains. In contrast to the reference cutin, the scattering patterns of cutins extracted by either ionic liquid show diffraction peaks indicating the presence of a crystalline component. The first diffraction peak at *q* =1.52 Å^−1^ and the second peak at *q* = 1.69 Å^−1^, noticeable for the cutin extracted by cholinium hexanoate for 170 h (Fig. 4A, blue curve), can be assigned to an orthorhombic crystal structure (space group Pnma, Miller indexes 110 and 200, respectively) commonly formed by compounds comprised of alkane-like chains such as triacylglycerols (b’ phase) (Mykhaylyk et al., 2007) or polyethylene (Bunn, 1944; Southern et al., 1972). This suggests that the extraction of cutin by the ionic liquids enriches this material with a crystalline component where some acyl chains tend to form crystals. A low level of branching of acyl chains in cutin possibly is favourable for the formation of an orthorhombic unit cell, which is thermodynamically more stable than the rotator phase formed by distorted alkane chains packed in a hexagonal array (Small, 1984). This observation is consistent with the DSC measurements (Fig. 4A) indicating that the cutin extracted by cholinium hexanoate for 170 h, containing the highest fraction of the acyl crystalline component, has the highest peak melting point. The third diffraction peak at *q* = 2.46 Å^−1^, observed for cutin extracted by cholinium hexanoate (Fig. 4B, purple and blue curves), cannot be related to the acyl chain crystalline structure. Its position is significantly shifted from a possible 020 peak at *q* = 2.55 Å^−1^ generated by the orthorhombic structure (Fig. 4B). The third peak is likely to be associated with a crystalline cellulose and can be assigned to 004 reflection of monoclinic cellulose I_β_ (space group P12_1_1) (Rongpipi et al., 2019). It has to be noted that the most intense 200 diffraction peak of the cellulose I_β_ expected at *q* = 1.63 Å^−1^ is not visible because of an overlap with the intense broad peak corresponding to the amorphous structure. Crystalline cellulose usually coexists with amorphous cellulose (Rongpipi et al., 2019). However, it would be difficult, if possible at all, to identify the cellulose amorphous component with its expected peak maximum intensity at *q* = 1.52 Å^−1^ from the broad diffuse peak observed by WAXS. Neither enzymatically-extracted cutin (reference cutin) nor cutin extracted by BMIM acetate reveal the presence of crystalline cellulose in their scattering patterns (Fig. 4B, black, cyan and grey curves), indicating that both extraction methods led to its successful removal. This observation together with the higher relative contribution of the oxygenated aliphatics region for the cutin extracted with BMIM acetate compared with the reference cutin (Table 1) suggests that some oxygenated aliphatics are lost during the enzymatic treatment.

**Fig. 4.**
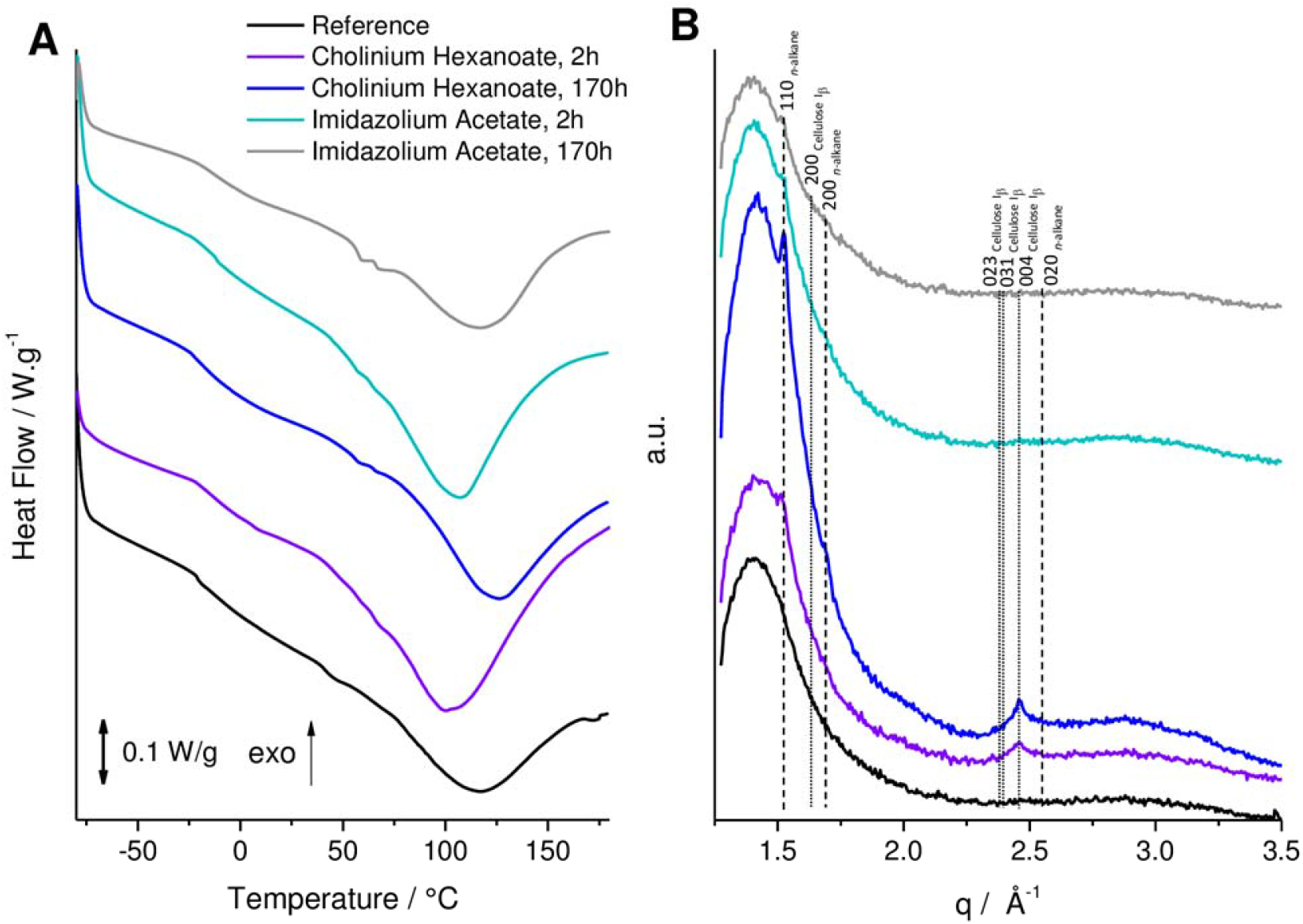
DSC thermograms (A) and WAXS patterns (B) collected for a reference enzymatically-extracted cutin (black curve) and cutin powders extracted from tomato peels using ionic liquids for various durations of the treatment [cholinium hexanoate (purple curve for 2 hours and blue curve for 170 hours) and imidazolium acetate (cyan curve for 2 hours and grey curve for 170 hours)]. The vertical straight lines in WAXS patterns indicate position of diffraction peaks of cellulose (dotted lines) and crystallised *n*-alkane chains (dashed lines). Miller indexes assigned to the lines correspond to cellulose I_β_ (monoclinic space group P12_1_1) and *n*-alkane chain packing (orthorhombic space group Pnma).

Finally, the relative abundances of hydrolysable cutin constituents were determined by GC-MS to disclose how the ionic liquid extraction methods influence the composition of the biopolymer compared to the reference method (Table 2, Supplementary Fig. S3). The reference cutin comprises *ca.* 13% of non-hydrolysable constituents, whereas those attained with an ionic liquid display significantly higher recalcitrance (*ca.* 20% to 30%), consistent with their estimated higher reticulation (Fig. 3B-C). In general, the monomeric compositions of the ionic liquid extracted cutins are similar to that of the cutin reference (and also to the starting material) (Table 2). Both the abundance and the diversity of fatty acids, decreased as the reaction time in the ionic liquid increased (Table 2). This was more pronounced when BMIM acetate was used for 170 hours, which rendered a cutin that is almost devoid of fatty acids and also containing nearly two times less dicarboxylic acids. The fatty acids carry a methyl end-group that is esterified to the biopolymer through a single bond. After 170 hours of reaction in BMIM acetate, the amount of 10,16-dihydroxyhexadecanoic acid that was lost from the biopolymer (*i.e.* solubilised) was nearly threefold higher than when cholinium hexanoate was used (data not shown).

**Table 2.** GC-MS quantitative analysis of the hydrolysable constituents identified in cutin samples purified using either cholinium hexanoate or 1-butyl-3-methlylimidazolium acetate (2, 15 or 170 h reactions). The reference cutin and the feedstock (*i.e*. untreated peels) were also analysed for comparison. Results are given as % (wt) (*n*=3). The identification yields (wt %) and the mass of the non-hydrolysable fraction (recalcitrance, %) are indicated below.

| Compound name | Untreated feedstock | Reference cutin | Compound abundance % (wt) |  |  |  |  |  |
| --- | --- | --- | --- | --- | --- | --- | --- | --- |
|  |  |  | cholinium hexanoate |  |  | BMIM acetate |  |  |
|  |  |  | 2h | 15h | 170h | 2h | 15h | 170h |
| **FATTY ACIDS** | **2.60 ± 0.13** | **1.52 ± 0.24** | **1.27 ± 0.15** | **0.88 ± 0.13** | **0.67 ± 0.09** | **1.96 ± 0.2** | **0.74 ± 0.1** | **0.10 ± 0.06** |
| hexadecanoic acid | 1.25 ± 0.04 | 0.74 ± 0.05 | 0.64 ± 0.04 | 0.41 ± 0.11 | 0.33 ± 0.03 | 0.63 ± 0.05 | 0.42 ± 0.03 | 0.10 ± 0.06 |
| octadeca-9,12-dienoic acid | 0.43 ± 0.15 | 0.23 ± 0.05 | 0.20 ± 0.03 | 0.11 ± 0.01 |  | 0.74 ± 0.12 |  |  |
| octadec-9-enoic acid |  | 0.24 ± 0.06 | 0.11 ± 0.02 | 0.09 ± 0.04 |  | 0.18 ± 0.03 |  |  |
| octadecanoic acid | 0.91 ± 0.12 | 0.31 ± 0.10 | 0.09 ± 0.00 | 0.13 ± 0.05 | 0.12 ± 0.02 | 0.18 ± 0.04 | 0.13 ± 0.06 |  |
| 4-oxocyclohexane-1-carboxylic acid |  |  | 0.23 ± 0.08 | 0.14 ± 0.12 | 0.22 ± 0.06 | 0.23 ± 0.00 | 0.19 ± 0.02 |  |
| **DICARBOXYLIC ACIDS** | **10.89 ± 1.08** | **18.38 ± 0.44** | **18.64 ± 0.49** | **19.2 ± 2.16** | **18.64 ± 0.58** | **18.45 ± 1.19** | **19.3 ± 0.21** | **9.99 ± 2.05** |
| nonanedioic acid <sup>(a)</sup> |  | 0.43 ± 0.02 | 0.26 ± 0.02 | 0.26 ± 0.15 | 0.36 ± 0.01 | 0.27 ± 0.02 | 0.22 ± 0.04 |  |
| hexadecanedioic acid | 0.56 ± 0.02 | 1.51 ± 0.07 | 1.30 ± 0.07 | 1.52 ± 0.10 | 1.22 ± 0.08 | 1.47 ± 0.07 | 1.42 ± 0.15 | 0.46 ± 0.07 |
| 8/9-hydroxyhexadecanedioic acid <sup>(b)</sup> | 10.33 ± 1.06 | 16.43 ± 0.38 | 17.08 ± 0.55 | 17.43 ± 2.01 | 17.06 ± 0.50 | 16.7 ± 1.14 | 17.66 ± 0.20 | 9.53 ± 0.23 |
| **ω-HYDROXY ACIDS** | **11.13 ± 0.35** | **15.04 ± 0.31** | **18.42 ± 0.63** | **17.42 ± 2.22** | **17.69 ± 1.28** | **18.95 ± 0.4** | **18.49 ± 1.06** | **22.90 ± 1.68** |
| 16-hydroxyhexadecanoic acid | 5.38 ± 0.16 | 8.17 ± 0.21 | 8.20 ± 0.12 | 8.50 ± 0.65 | 8.32 ± 0.21 | 8.35 ± 0.23 | 8.48 ± 0.39 | 15.71 ± 0.68 |
| 16-hydroxy-10-oxohexadecanoic acid | 3.02 ± 0.28 | 2.29 ± 0.14 | 6.39 ± 0.52 | 6.21 ± 0.97 | 6.28 ± 0.61 | 6.81 ± 0.05 | 6.62 ± 0.39 | 1.80 ± 0.12 |
| 9, 10-epoxy-18-hydroxyoctadecanoic acid | 1.14 ± 0.24 | 1.93 ± 0.09 | 1.73 ± 0.07 | 1.63 ± 0.38 | 1.27 ± 0.32 | 1.67 ± 0.13 | 1.60 ± 0.14 | 2.38 ± 0.53 |
| 9, 10-epoxy-18-hydroxyoctadecenoic acid | 1.60 ± 0.15 | 2.65 ± 0.20 | 2.10 ± 0.25 | 1.62 ± 0.18 | 1.82 ± 0.55 | 2.11 ± 0.09 | 1.79 ± 0.18 | 3.01 ± 0.59 |
| **POLYHYDROXYACIDS** | **55.05 ± 2.01** | **65.06 ± 0.65** | **61.67 ± 0.28** | **62.49 ± 4.06** | **63.01 ± 1.92** | **60.64 ± 1.69** | **61.47 ± 1.28** | **67.01 ± 1.44** |
| 10,16-dihydroxyhexadecanoic acid <sup>(c)</sup> | 53.22 ± 2.23 | 63.24 ± 0.98 | 58.83 ± 0.42 | 59.68 ± 5.03 | 61.20 ± 2.16 | 58.12 ± 1.89 | 58.97 ± 1.55 | 63.12 ± 2.42 |
| 9,10,18-trihydroxyoctadecanoic acid | 1.00 ± 0.11 | 0.94 ± 0.06 | 1.11 ± 0.32 | 1.19 ± 0.67 | 0.53 ± 0.00 | 0.85 ± 0.11 | 0.90 ± 0.13 | 1.51 ± 0.45 |
| 9,10,18-trihydroxyoctadec-12-enoic acid | 0.84 ± 0.39 | 0.89 ± 0.36 | 1.73 ± 0.07 | 1.63 ± 0.38 | 1.27 ± 0.32 | 1.67 ± 0.13 | 1.60 ± 0.14 | 2.38 ± 0.53 |
| **STEROLS\*** | **20.33 ± 1.07** |  |  |  |  |  |  |  |
| **Identification Yield (%)** | **56.38 ± 2.46** | **50.7 ± 2.38** | **58.1 ± 0.88** | **59.34 ± 6.63** | **56.39 ± 1.20** | **58.02 ± 2.45** | **60.35 ± 3.46** | **82.15 ± 0.20** |
| **Recalcitrance (%)** | **22.14 ± 2.98** | **13.32 ± 1.39** | **28.3 ± 4.98** | **27.47 ± 0.58** | **33.07 ± 0.56** | **19.75 ± 0.36** | **20.21 ± 1.06** | **18.15 ± 2.48** |

(a) Overestimated, overlapped with an unknown compound; (b) With possible presence of unspecific isomers; (c) Major 10,16-diOH, minors 9,16 and 8,16-*di*-OH.
\*Identified triterpenoids and sterols: β-amyrin (5.69 ± 0.44); α-amyrin (4.44 ± 0.20); δ-amyrin (7.53 ± 1.48) and stigmasterol (2.68 ± 0.49).

### A snap-shot of the molecular structure of cutin purified by cholinium hexanoate reveals extant free hydroxyls and free acids

Our data made evident the potential of using short time reactions with either ionic liquid to recover from tomato peels a cutin continuum displaying esterification/reticulation levels and composition near to that found *in planta*. In addition, cholinium hexanoate presents several advantages compared to the BMIM acetate. It cleaves fewer esters bonds, rendering a more esterified biopolymer (Fig. 3d), and contrary to the BMIM acetate, it is also biocompatible and biodegradable (Petkovic et al., 2010).

Recently, we resolved the molecular structure of *in situ* suberin using solution state NMR, upon its solubilisation in heated DMSO directly from cork after four hours of cryogenic milling (Correia et al., 2020). This inspired us to apply cryogenic milling for the solubilisation of a cutin extracted with cholinium hexanoate after two hours. Solving cutin’s molecular structure would create conditions to look “inside” its backbone, specifically to its esterification arrangement. The GC-MS analyses disclosed only the composing hydrolysable constituents (Table 2) and the solid-state analyses – ^13^C MAS NMR, DSC and WAXS (Fig. 3 and 4) – revealed only the bulky chemical functionalities and properties of the purified cutins. Only after 10 hours of cryogenic milling the cutin was solubilised in DMSO, reflecting cutin’s much lower solubility compared to suberin. We analysed the impact of the cryogenic milling process, especially the occurrence of oxidation reactions inside the grinding jar due to possible condensation of oxygen at low temperatures. Elemental analysis of cutin before and after the cryogenic milling process, revealed that the relative percentage of the tested elements, including oxygen (Supplementary Table S2), were unaltered after the treatment. Therefore, despite this solubility drawback, for the first time, a solution state ^1^H NMR could be acquired with good resolution showing the presence of many overlapping signals (Fig. 5A); an archetypal feature observed in other complex multifunctional polymers (Lyerla, 1980). The relative abundances of aliphatics, CH/CH_2_-X oxygenated aliphatics and aromatics were estimated through the integration of the ^1^H-spectrum as 70%, 27% and 3%, respectively. The assignment of ^1^H chemical shifts for the constituent monomers was then achieved through a combination of ^1^H-^1^H (COSY) and ^1^H-^13^C (HSQC, HMBC) correlation experiments (Supplementary Fig. S4 to S7). Previous NMR-based data of tomato cutin were attained through solution state NMR analyses of oligomeric structures obtained by methanolysis of tomato peels (Graça and Lamosa, 2010) and through HR-MAS NMR analyses of the tomato cutin swelled in DMSO (Deshmukh et al., 2003). These studies provided important baseline information for the assignment of the spectrum of cutin extracted with cholinium hexanoate for two hours (Supplementary Table S2).

**Fig. 5.**
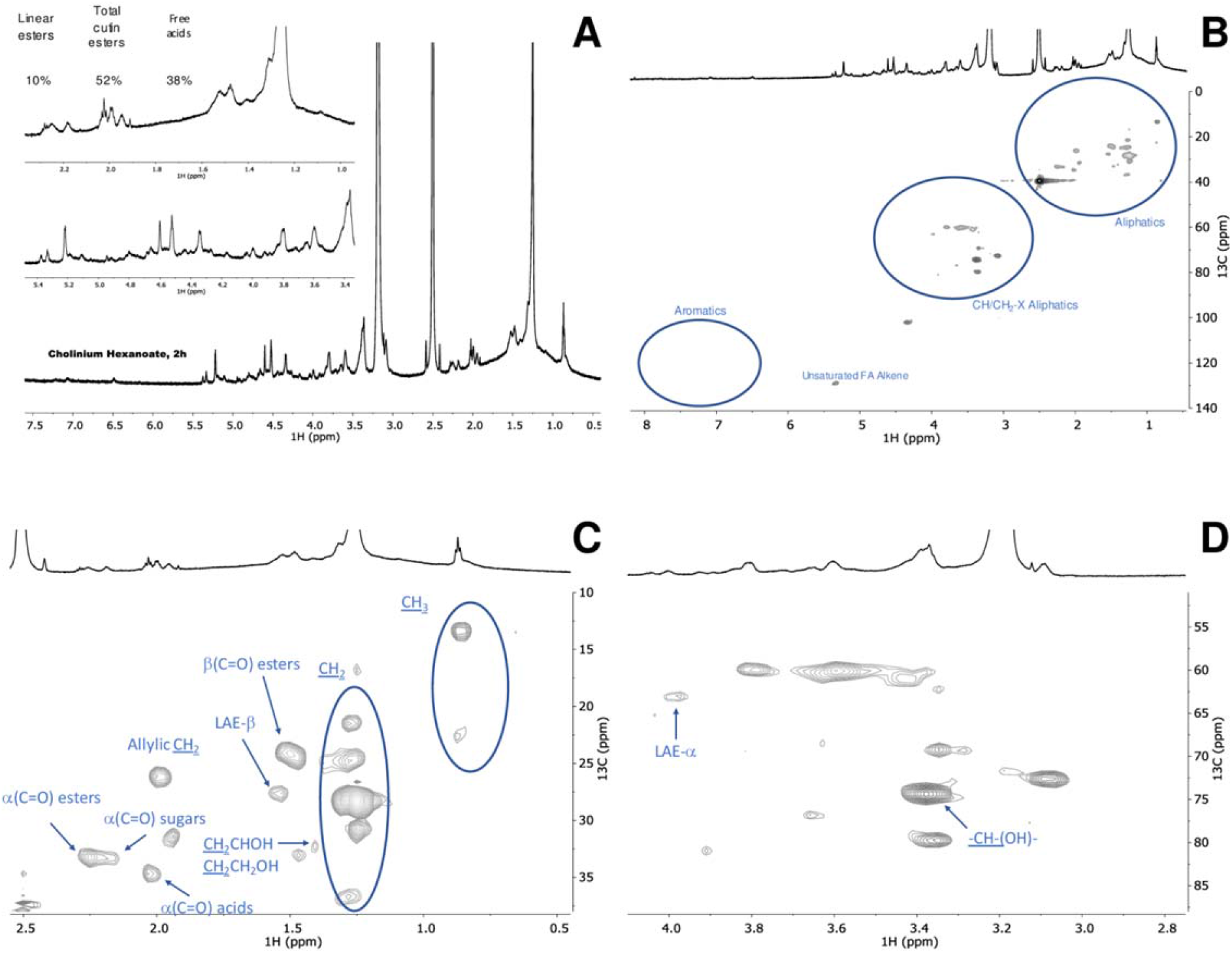
Wide-ranging NMR spectral characterisation of cutin isolated with cholinium hexanoate (2 h). The ^1^H NMR, with inserts focussing the aliphatic and oxygenated aliphatics regions (A); and the HSQC spectrum: full (B) and regions corresponding to aliphatics (C) and CH/CH_2_-X aliphatics (D) of the purified cutin. Some correlations (unlabelled) are uncertain or unidentified.

The full range HSQC spectrum of cutin is depicted in Fig. 5B, highlighting the regions corresponding to aliphatics and CH/CH_2_-X aliphatics as well as aromatics. A detailed analysis of the HSQC spectrum of the two aliphatics regions with the assignment of CH_2_ and CH_3_ groups from the aliphatic chains, the ester bonds and the free mid-chain hydroxyl groups is shown in Fig. 5C-D. Only secondary free hydroxyl groups were visible in the HSQC spectrum (CHOH *mid*-chain), consistent with their presence in cutin as suggested before (Philippe et al., 2016). This observation is in agreement with the demonstration that cholinium hexanoate does not cleave primary esters bonds (Fig. 1). Here we assigned the β-(C=O) esters to a ^1^H shift of 1.49 ppm and ^13^C shift of 24 ppm but we could not detect the signal of β-(C=O) acids, regardless that they have been assigned before in tomato cutin using HR-MAS NMR (Deshmukh et al., 2003). Deshmukh *et al.* (2003) assigned the signals of aliphatic esters, primary and secondary alcohols, free acids and α-branched carboxylic acids, yet the last two assignments could not be confirmed by HMBC. In the present study, the signals of the β-(C=O) acids possibly overlap with that of the esters and their differentiation from the small chemical shift differences observed in the acquired HSQC is virtually impossible. The signal α-(C=O) display two ^1^H signals with a ^13^C shift of 33 ppm, namely at 2.25 ppm and 2.19 ppm, which can be assigned to esters and acids, respectively. The α-(C=O) signal with a ^1^H shift of 2.17 ppm has been previously assigned to xylan esters (Zhang et al., 2016). Based on the detection of vestigial amounts of microcrystalline cellulose in the cutin extracted with cholinium hexanoate for 2 hours (Fig. 4B), this signal may be associated to the presence of cellulose esters. Analysis of the cutin extracted with BMIM acetate (upon its cryogenic milling), which is apparently devoid of microcrystalline cellulose (Fig. 4B), showed that the signal α-(C=O) display a ^13^C shift of 33 ppm and only a ^1^H shift of 2.26 ppm (Supplementary Fig. S6). Finally, to precisely assign the free acids in the cutin extracted with cholinium hexanoate for 2 hours, we acquired the HMBC spectrum that confirmed their signal at a ^13^C shift of 35 ppm and a ^1^H shift of 2.02 ppm (Supplementary Fig. S7). This observation is consistent with that previously assigned in cork suberin where the signal of the acid is at a ^13^C shift of 36 ppm and a ^1^H shift of 2.03 ppm, and that of the esters displays a ^13^C shift of 34 ppm and a ^1^H broad shift from 2.33-2.27 ppm (Correia et al., 2020).

Based on the assignments defined above, we calculated through integration of the signals in the ^1^H NMR the relative abundance of free acids, of total esters (comprising primary and secondary aliphatic esters yet excluding sugar esters) and of linear esters as 38%, 52% and 10%, respectively (Fig. 5A, *see text-insert*). No acylglycerol bonds were detected in the HSQC analyses of cutin (Fig. 5B), consistent with the very low abundance of glycerol in tomato cutin (Fich et al., 2016). We hypothesise that the free acids detected in the cutin spectra might mostly account for their natural occurrence, though one cannot exclude, at this stage, that some aliphatic esters might underwent cleavage in the presence of cholinium hexanoate.

### Ionic liquid extraction followed by solution NMR as a new tool to scrutinise the impact of specific mutations in the molecular structure of cutin from Micro-Tom tomatoes

Solving the molecular structure in solution of a near native cutin isolated from a processing tomato cultivar, challenged us to test the suitability of the established cholinium hexanoate extraction for two hours, for the purification and systematic characterisation of cutins isolated from the tomato miniature cultivar Micro-Tom particularly well-suited for laboratory studies (Just et al., 2013; Garcia et al., 2016). To introduce known diversity in native cutin composition and structure, we further used both the wild type and the *gpat6* (*GLYCEROL-3-PHOSPHATE ACYLTRANSFERASE* gene) and the *cus1* (*CUTIN SYNTHASE* gene) (Petit et al., 2016; Philippe et al., 2016; Petit et al., 2017) mutants that show phenotypes with altered cutin composition and altered degree of intra-chain branching. In particular, in the *gpat6* mutant (formerly named cutin-deficient mutant *cud1*; (Philippe et al., 2016)) the synthesis of the major cutin precursor is hampered, hence a thinner cuticle is produced with overall decreased levels of cutin, which is enriched in fatty acids (Petit et al., 2016). In contrast, in the *cus1* tomato mutants, cutin polymerization is impaired (Girard et al., 2012; Yeats et al., 2012) and the esterification of secondary OH groups of the dihydroxy acids is significantly reduced (Philippe et al., 2016). To minimize any possible effect of the environmental conditions on the expression of the fruit cuticle phenotype, the *gpat6* and *cus1* mutants were grown side-by-side with wild type plants. The relative abundance of the hydrolysable constituents in the Micro-Tom cutins purified with cholinium hexanoate is depicted in Table 3. In general, the observed diversity/abundances of hydrolysable constituents are similar to that previously reported (Petit et al., 2016; Philippe et al., 2016), regardless of some variations, possibly due to disparities in tomato growth conditions in the greenhouse (season, light, temperature and hygrometry). In addition, the cutins which originated from the mutants show an increase in the relative abundance of non-hydrolysable constituents compared to the wild type (*ca.* 10% increase), and their identification yields decreased nearly 20% due to higher diversity of unidentified monomers (Table 3). Cutin from both mutants display higher relative abundance of fatty acids and dicarboxylic acids (nearly tenfold and twofold, respectively) and lower of ɷ-hydroxyacids (five to three times) compared to the wild type cutin.

**Table 3.** GC-MS quantitative analysis of the hydrolysable constituents in cutin samples purified using cholinium hexanoate (2 h reaction) from Micro-Tom tomatoes: wild type, *cus1* and *gpat6* plants. Results are given as % (wt) (*n*=3). The identification yields (wt %) and the mass of the non-hydrolysable fraction (recalcitrance, %) are indicated below.

| Compound name | Compound abundance % (wt) |  |  |
| --- | --- | --- | --- |
|  | cholinium hexanoate (2 h) |  |  |
|  | *wt* | *cus1* | *gpat6* |
| **FATTY ACIDS** | **0.50 ± 0.04** | **6.06 ± 0.81** | **5.60 ± 0.88** |
| hexadecanoic acid | 0.27 ± 0.02 | 2.09 ± 0.26 | 1.59 ± 0.23 |
| 9,12-octadecadienoic |  | 3.00 ± 0.59 | 2.22 ± 0.42 |
| 9-octadecenoic acid | 0.24 ± 0.06 |  | 1.20 ± 0.16 |
| octadecanoic acid |  | 0.96 ± 0.12 | 0.59 ± 0.11 |
| **DICARBOXYLIC ACIDS** | **4.68 ± 0.32** | **7.23 ± 0.55** | **7.92 ± 0.5** |
| nonanedioic acid <sup>(a)</sup> |  | 1.75 ± 0.37 | 0.88 ± 0.05 |
| hexadecandioic acid | 0.60 ± 0.10 |  | 1.95 ± 0.23 |
| 8/9-hydroxyhexadecanedioic acid <sup>(b)</sup> | 4.08 ± 0.23 | 5.48 ± 0.24 | 5.09 ± 0.25 |
| **ω-HYDROXY ACIDS** | **5.47 ± 0.30** | **1.19 ± 0.02** | **1.56 ± 0.08** |
| 16-hydroxyhexadecanoic acid | 3.97 ± 0.23 | 1.19 ± 0.02 | 1.12 ± 0.09 |
| 16-hydroxy-10-oxohexadecanoic acid | 0.71 ± 0.04 |  | 0.44 ± 0.02 |
| 9, 10-epoxy-18-hydroxyoctadecanoic acid | 0.25 ± 0.05 |  |  |
| 9, 10-epoxy-18-hydroxyoctadecenoic acid | 0.54 ± 0.02 |  |  |
| **POLYHYDROXYACIDS** | **89.35 ± 0.63** | **85.33 ± 1.32** | **84.86 ± 1.46** |
| dihydroxyhexadecanoic acid <sup>(c)</sup> | 87.91 ± 0.48 | 83.87 ± 1.52 | 77.71 ± 2.50 |
| 9,10,18-trihydroxyoctadecanoic acid | 0.62 ± 0.06 | 0.91 ± 0.18 | 3.84 ± 0.76 |
| 9,10,18-trihydroxyoctadec-12-enoic acid | 0.82 ± 0.09 | 0.54 ± 0.04 | 3.31 ± 0.49 |
| **STEROLS\*** |  | **0.19 ± 0.04** |  |
| **Identification Yield (%)** | 57.99 ± 1.26 | 37.49 ± 0.74 | 36.05 ± 1.75 |
| **Recalcitrance (%)** | 32.6 ± 3.68 | 44.26 ± 2.96 | 41.28 ± 0.97 |

(a) Overestimated, overlapped with an unknown compound; (b) With possible presence of unspecific isomers; (c) Major 10,16-diOH, minors 9,16 and 8,16-*di*-OH. \*Identified sterols: stigmasterol

To confirm that free acids naturally occur in cutin (hence differentiating these from free acid groups formed during the ionic liquid extraction), we compared the spectrum of cutin from the wild-type cultivar purified by the ionic liquid with that of the solubilised cuticle *via* cryogenic milling (Fig. 6). The obtained ^1^H NMR (Fig. 6A-B) and HSQC spectra (Fig. 6 C-F) are very similar, regardless that the presence of non-cutin constituents in the cuticle contributes to the appearance of many new signals, yet to be assigned, *e.g.* in the CH_3_ region (Fig. 6D). Importantly, the signals previously assigned to free acids - α-(C=O) acids - are visible in both samples (Fig. 6C-D), which were confirmed in the corresponding HMBC spectra (Supplementary Fig. S10). Accordingly, the free acids detected in the ionic liquid purified cutins (Fig. 5A-C and 6A-B) reflect their natural presence. The signal attributed to the terminal hydroxyls was only detected in the spectrum of the ionic liquid extracted Micro-Tom cutin (Fig. 6E). This observation suggests that the cholinium hexanoate treatment cleaved some primary esters in the Micro-Tom cutin, contrary to that observed for the cutin derived from the peels of processing tomatoes (Fig. 5D). One possibility is that the cleavage of primary esters is greatly influenced by the native arrangement of the polymer.

**Fig. 6.**
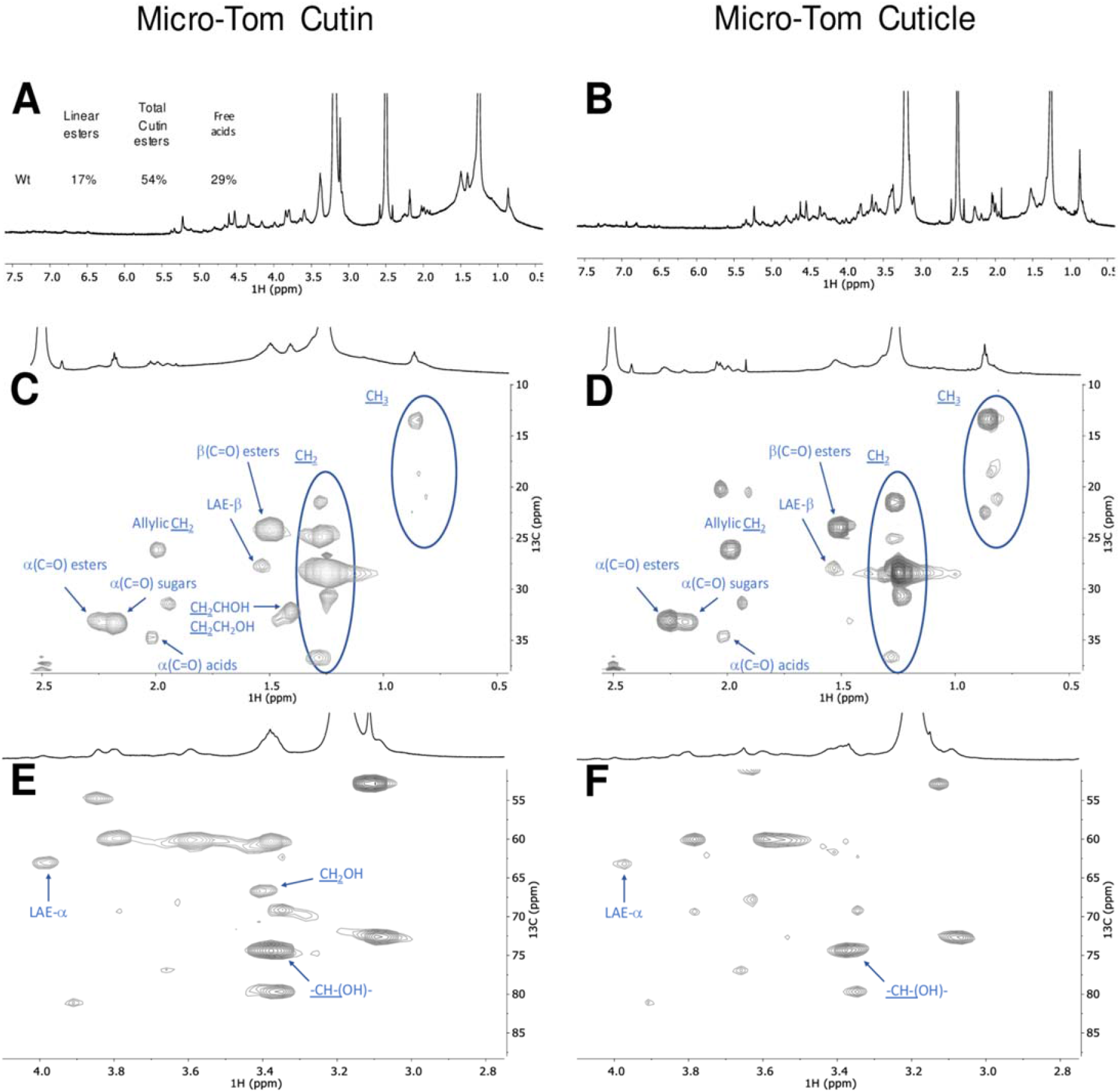
Wide-ranging NMR spectral characterisation of Micro-Tom cutin isolated with cholinium hexanoate (2 h) and Micro-Tom untreated cuticle. The ^1^H NMR spectra of both samples (A - cutin and B - cuticle), where the text-inserts indicate the relative abundance (%) of Linear aliphatic esters (LAE-α), total esters (α(C=O) esters) and the free acids (α(C=O) acids); HSQC regions corresponding to aliphatics (C - cutin and D - cuticle) and CH/CH_2_-X aliphatics (E - cutin and F - cuticle). Some correlations (unlabelled) are uncertain or unidentified.

The impact of the mutations is seen by the relative abundances of aliphatics, CH/CH_2_-X oxygenated aliphatics, and aromatics, in the ^1^H-spectra which were estimated as 71%, 29% and for the wild-type (Fig. 6A), as 46%, 50% and 4% for *gpat6* mutant (Fig. 7A), and as 39%, 59% and 2% for the *cus1* mutant (Fig. 7B), respectively. Contrary to the wild type, in both mutants the signal assigned to free acids could not be detected (Fig. 7) (Fig. 6A, *see text-inserts*). To confirm this observation, we compared the spectrum of the cutin from the *cus1* mutant purified by the ionic liquid with that of the solubilised *cus1* cuticle *via* cryogenic milling (Supplementary Fig. S16). The obtained HSQC spectra confirmed the absence of free acids in this mutant, furthering that the observed absence of this chemical group in the *cus1* and *gpat6* cutins is a consequence of the mutations and not of the sample processing.

**Fig. 7.**
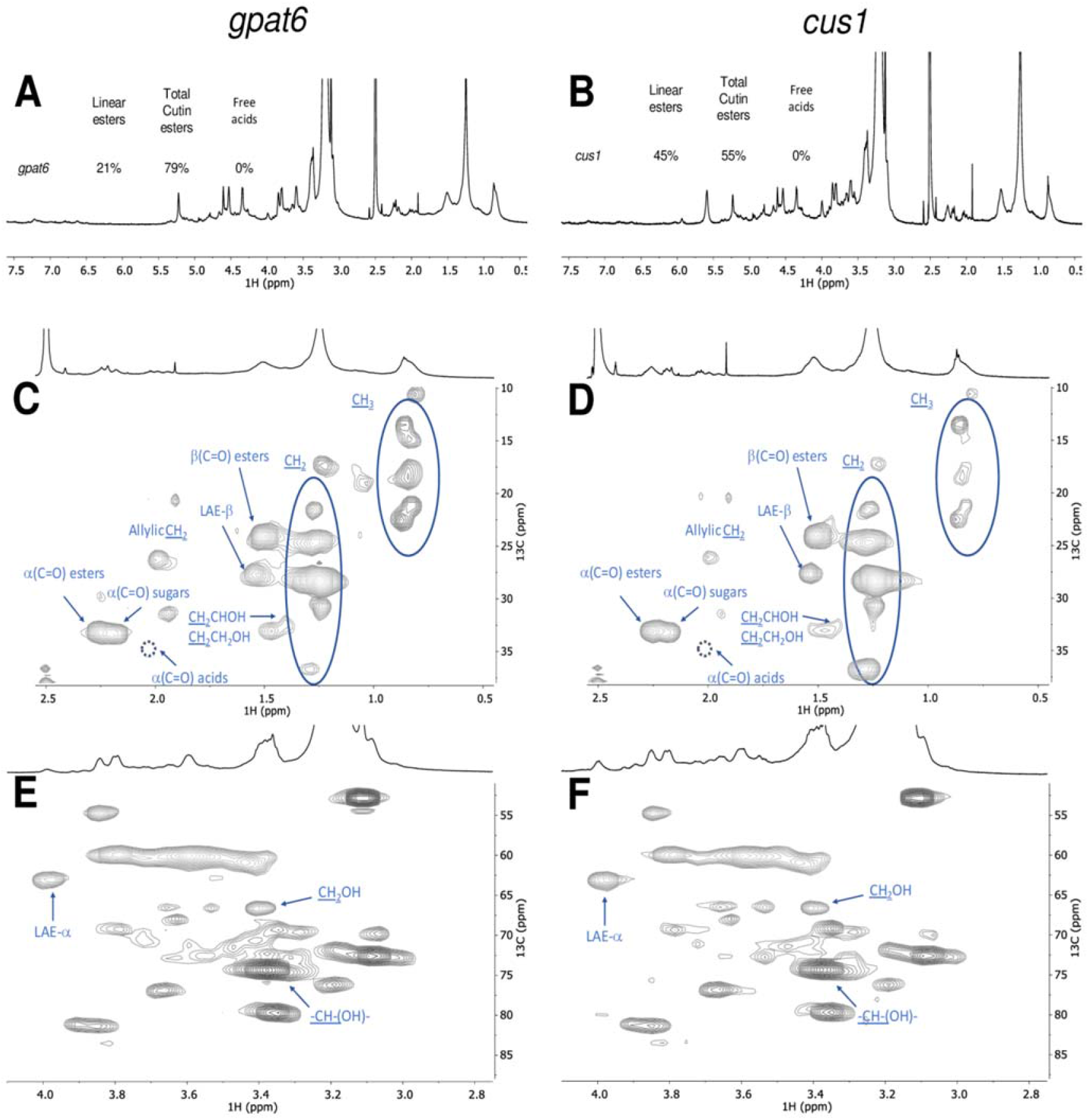
Wide-ranging NMR spectral characterisation of Micro-Tom cutins isolated with cholinium hexanoate (2 h) from the *cus1* and *gpat6* mutants. The ^1^H NMR spectra of both samples (A - *cus1* - and B - *gpat6*), where the text-inserts indicate the relative abundance (%) of Linear aliphatic esters (LAE-α), total esters (α(C=O) esters) and the free acids (α(C=O) acids); HSQC regions corresponding to aliphatics (C - *cus1* and D - *gpat6*) and CH/CH_2_-X aliphatics (E - *cus1* and F - *gpat6*). Some correlations (unlabelled) are uncertain or unidentified. The absence of the signal assigned to α(C=O) acids is marked by a dashed circle. For simplicity the wide-ranging NMR spectrum of the untreated cuticle from the *cus1* mutant is not shown (detailed in Supplementary Fig. S16).

Based on the ^1^H-spectral information, we also estimated the relative abundance of aliphatic esters (total) and primary aliphatic esters in the cutin of the mutants (Fig. 7, *see text-inserts*). Accordingly, the ratio of total esters *versus* linear esters is comparable in the wild type and the *gpat6* mutant but significantly lower in the *cus1* mutant. By other words, compared to the wild type, the *gpat6* mutant show similar amount of linear esters and of secondary esters, contrary to the *cus1* mutant that shows more than a twofold increase in linear esters but also the lowest esterification level (*i.e.* amount of secondary esters) (Fig. 7A-B, *see text-insert*).

The magnification of the HSQC regions corresponding to aliphatics and CH/CH2-X aliphatics for the cutins of the mutants is also shown (Fig. 7C-F). For both mutants, the signals assigned to terminal hydroxyls are visible (Fig. 7E-F), similar to that observed in the wild type cutin (Fig. 6E). The detected CH_3_ groups are apparently enriched in both mutants compared to the wild type, consistent with the observed increase in the relative abundance of hydrolysable fatty acids for the mutants (Table 3) and with that reported before (Petit et al., 2016; Philippe et al., 2016). No acylglycerol was visible in the HSQC spectra of the cutin from the mutants, possibly as their abundance are below the detection limits of the analytical technique. The mutants show more non-assigned signals compared to the wild-type cutin, consistent with the observed lower identification yields in the GC-MS (Table 3). This might reflect also an altered diversity of cuticular-polysaccharides in the mutants; one hypothesis that deserves further analysis in the near future and is sustained by the recently published results on *cus1* mutant (Philippe et al., 2020).

## Discussion

Considerable advances have been made in the recent years on cuticle formation and properties (Nawrath et al., 2013). However, while the successive steps of cutin biosynthesis, transport of precursors and polymerisation in the epidermal cell walls begin to be well untangled (Fich et al., 2016), the questions of the fine structure of the cutin layer and its association with polysaccharides are still largely unresolved (Philippe et al., 2020). An example of the intricate relationships between cutin, cell walls and resistance to pathogens is that provided by the tomato *gpat6* mutant analysed herein in which the mutation has a profound impact not only on cutin synthesis (Petit et al., 2016) but also on cell walls (Philippe et al., 2020) and resistance to filamentous pathogens (Fawke et al., 2019). New insights into the structure and composition of native *gpat6* cutin would considerably help deciphering the underlying mechanisms. More generally, the simple and rapid cutin extraction method described here, which preserves cutin in a near native state, will help our understanding of the role of cuticle in plant evolution and diversity (Yeats et al., 2012; Yeats et al., 2014), plant development (Ingram and Nawrath, 2017), mechanical properties of the organ surface (España et al., 2014; Mazurek et al., 2017), resistance to pathogens (Chassot et al., 2007) and fruit quality (Petit et al., 2017).

### Advantages of ionic liquid extraction with respect to conventional cutin extraction methods

Our ionic liquid cutin extraction method performed on tomato peels demonstrates that subcuticular polysaccharides (which were found in the filtrate) are removed similar to that of the enzyme treatment but in a considerable shorter period of time (*i.e.* 2 h instead of days, and even weeks) (Chatterjee et al., 2012). Also, the ionic liquid extraction does not require any specific dewaxing step. When extracted with either ionic liquid, a cutin-continuum is isolated, strengthening that the ionic liquid does not impact significantly the cutin polyester. This opposes to that previously reported by us for suberin extraction from cork using cholinium hexanoate where nanoparticles of the biopolymer are isolated (Correia et al., 2020) since the ionic liquid catalyses the mild cleavage of acylglycerol ester bonds (Ferreira et al., 2014) (which are not representative in tomato cutin).

### Cholinium hexanoate extraction preserves features of cuticular polysaccharides

In the present study, the purification of the cutin-continuum by the ionic liquids is essentially due to the dissolution of the subcuticular polysaccharides and at a minor extent to ester cleavage. Under the conditions of the extraction here used, both ionic liquids cleaved very inefficiently the esters bonds; however, cholinium hexanoate apparently cleaves only the acylglycerol bonds, whereas BMIM acetate can cleave also linear esters bonds. Hence, the cholinium hexanoate presents the advantage of biocompatibility and of a milder cleavage of the polymer backbone. Their most remarkable difference is that only the cholinium hexanoate could preserve features of cuticular polysaccharides, which were specifically associated to the presence of microcrystalline cellulose. Cellulose with high levels of crystallinity was recently identified within the group of cutin embedded polysaccharides (Philippe et al., 2020), consistent with the ability of the used ionic liquid process for a speedy recovery of cutin with a near native structure. This opens unexpected possibilities for exploiting both ionic liquids, alone or in combination, to study the association and function of cuticular polysaccharides.

### Cholinium hexanoate extraction confirms the presence of free hydroxyls in native cutin and highlights differences in free fatty acid composition between wild type and mutants

Finally, the solution NMR spectra of cutins purified by cholinium hexanoate and of the matching cuticles (both of which solubilised with the aid of cryogenic milling) strengthened that some free hydroxyls exist *in situ*, consistent with that previously reported by others (Petit et al., 2016; Philippe et al., 2016). Results from a systematic NMR analysis of cutins purified by cholinium hexanoate from both wild-type and mutants of the same genotype (Micro-Tom) show increased diversity of fatty acids in both mutants yet only the *cus1* mutant show a significantly reduced esterification degree. These results are consistent with the function of the enzymes inactivated: *cus1* is a polymerizing enzyme (Girard et al., 2012; Yeats et al., 2012) while *gpat6* catalyses the synthesis of cutin precursors (Petit et al., 2016). Remarkably, these results are also consistent with already published information on the mutants, which were obtained through a totally independent approach (Philippe et al., 2016). *In situ* analysis of cutin esterification levels by benzyl etherification of enzyme-treated cutin from tomato fruit peel showed that all/midchain hydroxylation of dihydroxy acids is increased strongly in *cus1* (linear polymers) but remained unaffected in *gpat6* mutant, as in wild type (normal inter-branching).

Remarkably, the NMR results of the ionic liquid extracted cutins are strongly suggestive that naturally occurring free acids exists in the wild-type tomatoes (detected also in the cuticle) however lacking in the mutants (lacking also in their cuticles, as observed for the *cus1* cuticle). This might be due to the thinner mutant cuticles where a total esterification of the cutin monomers could be more easily achieved. This deserves a detailed analysis in the near future, especially as: (i) we could not yet detect the signal assigned to β(C=O) acid, only that of the α(C=O) acids that was assigned through HMBC spectrum, and (ii) the signal assigned to the α(C=O) esters partially overlaps with that of the α(C=O) sugars in the analysed tomato cutins.

### Conclusions

The proof-of-concept of the efficiency and reliability of the ionic liquid cutin extraction method described here was done using tomato peel as a model. Because of its simplicity, this method should be broadly applicable to other tissues and to other plant model and crop species, as confirmed by preliminary experiments. In the near future quantitative methods (and better spectral resolution for solving yet unknown signals) will require development in order to understand better how cutin molecular structure (and its association with cuticular polysaccharides) is impacted by mutations or along the development of the plant. In addition, our study emphasises the suitability of exploiting ionic liquid extractions as an easy and scalable approach for exploiting plant lipid polymers as a bio-resource for a diversity of applications. The ionic liquid processes here tested can be systematically tuned, *e.g.* time, temperature, composing ions, to ensure recovery of a cutin with different degrees of structural preservation. Finally, the solution NMR methodologies developed here constitute now essential tools to fingerprint the multi-functionality and the structure of cutin *in planta*. Based on all the analyses done on the polymer morphology, thermal properties and chemistry, we are confident that short-time reactions with cholinium hexanoate can ensure the isolation of cutin carrying minimal disruption of its polymeric network – yielding the closest to a native configuration reported to date.

## Material and Methods

### Plant Material

Peels from the processing tomato (*Solanum lycopersicum* ‘Roma’) were manually removed, thoroughly washed and then dried until constant weight at 60 °C. After drying, the peels were milled using a Retsch ZM200 electric grinder (granulometry 0.5 mm; 10000 rpm) and stored at room temperature for further processing. Micro-Tom cultivar tomatoes from both wild type and mutants plants (which were generated by an ethyl methanesulfonate (EMS) mutagenesis (Just et al., 2013)) were cultivated as previously reported (Rothan et al., 2016), and processed as described above. All tomato fruits used were in the red ripe developmental stage.

### Chemicals

1-butyl-3-methyl-imidazolium acetate (>98%) was purchased from io-li-tec; sodium hydroxide (>98%) from José Manuel Gomes dos Santos; methanol (≥99.8%), dimethyl sulfoxide (DMSO, >99.99%), hexane (>95%), chloroform (>99.98%), dichloromethane (>99.99%) from Fisher Chemical; cholinium hydrogen carbonate (~80% in water), hexanoic acid (>99.5%), sodium azide (≥99.5%), sodium acetate (≥99%), cellulase (*Aspergillus niger*) and pectinase (*Aspergillus aculeatus*) from Sigma Aldrich. Cholinium hexanoate was synthesised by dropwise addition of hexanoic acid to aqueous cholinium hydrogen carbonate in equimolar quantities, as previously described (Petkovic et al., 2010).

### Ionic liquid hydrolysis of standard fatty acids

Glyceryl trioctanoate and octyl octanoate (*ca*. 50 mg) were mixed with either ionic liquid (1:10 ratio) at 100 °C, without stirring, during 2, 6 or 24 hours. At the end of the reaction, the mixture was rapidly cooled to room temperature in ice, acidified to pH 3/3.5 with 1 M HCl solution, spiked with a known concentration of heptadecanoic acid (internal standard), and extracted three times using dichloromethane/water partition. The dried combined organic phases were derivatised with N,O-bis(trimethylsilyl)trifluorocetamide in pyridine (5:1), during 30 min at 90 °C. The TMS derivatives in the organic fractions were then analysed by GC-MS as previously described (Ferreira et al., 2014) with minor modifications (ramp temperature: 60 °C, 4 °C/min until 280 °C during 15 min, with source at 230 °C and electron impact ionization of 70 eV) (see equipment below). Triplicate independent reactions were performed.

### Cutin Extractions

#### Enzymatic Process

Cutin was isolated from tomato as previously described (Chatterjee et al., 2012). In brief, tomato peels were immersed in an enzymatic cocktail containing 4 ml of pectinase, 0.2 g of cellulase, 13 mg NaN_3_ and 196 ml of 50 mM sodium acetate buffer, and incubated at 31 °C for 24 hours with constant shaking. The isolated cuticles were successively dewaxed during 36 hours by Soxhlet extraction with methanol, chloroform and hexane (1:1:1), finally freeze dried and stored at room temperature. This cutin is used as a reference material in the present study.

#### Ionic liquid Process

Cutin was extracted from tomato peels as previously described for the extraction of suberin from cork (Ferreira et al., 2014), with slight modifications. In brief, 2 g of tomato peel powder were mixed in 20 g cholinium hexanoate or BMIM acetate and incubated for a defined period of time (100 °C, without stirring). The reaction was stopped by the addition of 160 mL of DMSO. The polymer was recovered by filtration using a nylon membrane filter (0,45 μm); then washed with an excess of deionized water with the aid of centrifugation (Eppendorf 5804 R centrifuge, 5000 rpm at 4 °C for 30 minutes).

### Treatment of cutin extracted using cholinium hexanoate (2h) with Trifluoroacetic Acid

600 mg of the cutin obtained through extraction with cholinium hexanoate for 2 h were treated with 1 M of aqueous TFA solution during 60 minutes at 110 °C. The reaction mixture was filtered, and the insoluble material was washed with stirring using chloroform-methanol (1:1 v/v) for 2 h. The organic-insoluble material was separated by filtration, freeze-dried, and analysed by ^13^C MAS NMR.

### Microscopic analyses

Scanning electron microscopy (SEM) (microscope JEOL JSM-7001F) was used to analyse the cutin samples.

### Cryogenic grinding process

A RESTCH Cryomill equipped with a 25 mL grinding jar with 6 zirconium oxide grinding balls (10 mm) was used. To optimise the solubilisation level of cutin needed for attaining high NMR spectral resolution, cutin samples were cryogenically milled at −196 °C (liquid nitrogen) as follows: 3 min of pre-cooling followed by 9 milling cycles, each comprising 3 min of milling at 30 Hz plus 0.5 min of intermediate cooling at 5 Hz. The ensuing samples were analysed by ^1^H NMR (3 mg dissolved in 400 μL of DMSO-*d*_6_) and the 200 milling cycles were selected for systematically process cutin samples prior to their 2D NMR analysis.

### Nuclear Magnetic Resonance (NMR) analyses

Solution state NMR spectra were recorded using an Avance II + 800 MHz (Bruker Biospin, Rheinstetten, Germany) spectrometers, with exception of ^1^H-^13^C HMBC spectra that were acquired using an Avance III 800 CRYO (Bruker Biospin, Rheinstetten, Germany). All NMR spectra (^1^H, ^1^H-^1^H COSY, ^1^H-^13^C HSQC) were acquired in DMSO-*d*_6_ using 5 mm diameter NMR tubes, at 60 °C as follows: 3 mg of cryomilled cutin in 400 μL of DMSO-*d*_6_. ^13^C Magic Angle Spinning Nuclear Magnetic Resonance (^13^C MAS NMR) spectra were acquired on cutin samples (± 250 mg) were packed into 7 mm o.d. zirconia rotors (after grinded if needed), equipped with Kel-F caps. ^13^C MAS with High-Power CW Decoupling spectra were obtained at 75.49 MHz, on a Tecmag Redstone/Bruker 300WB, with spinning rates of 3.1-3.3 kHz. In these experiences 90° RF pulses of around 4.5 μs and relaxation delays of 3 s were used. ^13^C chemical shifts were referenced with respect to external glycine (^13^CO observed at 176.03 ppm). MestReNova, Version 11.04-18998 (Mestrelab Research, S.L.) was used to process the raw data acquired in the Bruker spectrometers.

### Differential scanning calorimetry (DSC)

Calorimetric analyses were carried out in a TA Instruments Q200 calorimeter connected to a cooling system and calibrated with different standards (indium, empty cap). The sample weights ranged from 9 to 11 mg. A temperature interval from −80 °C to 220 °C has been studied and the used heating/cooling rate was 10 °C·min^−1^.

### Wide-Angle X-ray Scattering (WAXS)

WAXS data were collected using a laboratory SAXS/WAXS beamline (Xeuss 2.0, Xenocs, Grenoble, France) equipped with a liquid gallium MetalJet X-ray source (Excillum, Sweden) (wavelength *λ* = 1.34 Å), FOX 3D Ga single reflection X-ray mirror and two two sets of motorized scatterless slits for beam collimation, and a Pilatus 100k two-dimensional (2D) pixel WAXS detector (Dectris, Switzerland). Loose cutin powder samples were enclosed between two flat kapton films and mounted on the beamline sample stage (the total sample thickness is about 1 mm). 2D WAXS patterns were recorded in a transmission mode over a *q* range of 1.3 Å^−1^ to 3.5 Å^−1^ [where *q* = (4*π*sin*θ*)/*λ* = 2*π*/*d* is the length of the scattering vector, *θ* is one-half of the scattering angle and *d* is spacing in real space] using exposure time of 1200 seconds. The WAXS data were reduced (calibrated, integrated and background-subtracted) using the Foxtrot software package supplied with the instrument.

### Gas Chromatography-Mass Spectrometry (GC-MS)

An Agilent gas chromatograph (7820A) equipped with an Agilent (5977B) mass spectrometer (quadrupole) was used. First, to release the hydrolysable constituents, the samples were treated with a solution of 0.5 M NaOH in methanol/water (1:1, v/v) at 95 °C, during 4 hours; cooled to room temperature and acidified to pH 3/3.5 with 1 M HCl, then extracted by dichloromethane/water partition (5X). The non-hydrolysable fraction was recovered by filtration (0.2 μm, nylon filters), washed, freeze-dried, and weighted (recalcitrance). The dried combined organic extracts were sequentially derivatised (30 min, 90 °C): firstly, 2.0 M (trimethylsilyl)diazomethane in hexane, mixed in a methanol:toluene 2.5:1 solution (3:2); and secondly, N,O-bis(trimethylsilyl)trifluoroacetamide containing 1% of trimethylchlorosilane in pyridine (5:1). The derivatives were then analysed by GC-MS (HP-5MS column) with the following ramp temperature: 80 °C, 4 °C/min until 310 °C during 15 min. MS scan mode, with source at 230 °C and electron impact ionization (EI+, 70 eV) was used for all samples. The GC-MS was first calibrated with pure reference compounds (representative monomers: heptadecanoic acid, hexadecanedioic acid and ferulic acid) relative to hexadecane (internal standard). Each sample was analysed in triplicates. Data acquisition was accomplished by MSD ChemStation (Agilent Technologies); compounds were identified based on the equipment spectral library (Wiley-NIST) and references relying on diagnostic ions distinctive of each derivative and its spectrum profile (Supplementary Table S1).

## Acknowledgments

We acknowledge funding from the European Research Council through grant ERC 2014-CoG-647928, from the European Union’s Horizon 2020 research and innovation programme within the project 713475 – FLIPT – H2020-FETOPEN-2014-2015 and from Fundação para a Ciência e Tecnologia (FCT) through the grant UID/Multi/04551/2019 (Research unit GREEN-it “Bioresources 4 Sustainability”) and the projects PTDC/AGR-TEC/1191/2014AAC. The solution NMR data was acquired at CERMAX, ITQB-NOVA, Oeiras, Portugal with equipment funded by FCT. C.J.S.M. is grateful to Aralab, Portugal, for the PhD contract 06/PlantsLife/2017. O.O.M. thanks EPSRC for a capital equipment grant to purchase the Xenocs/Excillum SAXS/WAXS laboratory beamline (EP/M028437/1). The authors are thankful to Manolis Matzapetakis (ITQB NOVA) and to Maria João Ferreira (IST) for support in the solid-state NMR analyses, and to Pedro Lamosa and Maria C. Leitão (ITQB NOVA) for support in the solution NMR and chromatographic analyses, respectively. Finally, we are extremely grateful to Bénédicte Bakan and Didier Marion for fruitful scientific discussions during the manuscript preparation.

